# ProxLogs: Miniaturised proximity loggers for monitoring association behaviour in small mammals

**DOI:** 10.1101/2021.02.28.432842

**Authors:** Lucinda Kirkpatrick, Ivan Hererra Olivares, Apia Massawe, Christopher Sabuni, Herwig Leirs, Rafael Berkvens, Maarten Weyn

**Affiliations:** Evolutionary Ecology, Universiteit Antwerpen, Antwerp, Belgium; IDLab, Faculty of Applied Engineering, Universiteit Antwerpen–imec, Antwerp, Belgium; Pest Management Centre, Sokoine University of Agriculture, Morogoro, Tanzania

**Keywords:** Small animals, Open Source, Contact network, Proximity loggers, disease transmission, association behaviour, biologging, Bluetooth Low Energy

## Abstract

1. The ability to monitor associations between wild animals is essential for understanding the processes governing gene transfer, information transfer, competition, predation and disease transmission.
2. Until recently, such insights have been confined to large, visible or captive animals. However, the rapid development of miniature sensors for consumer electronics is allowing ecologists to monitor the natural world in ways previously considered impossible.
3. Here we describe miniature (<1g) proximity loggers we have developed that use Bluetooth Low Energy transmission to register contacts between individuals. Our loggers are open source, low cost, rechargeable, able to store up to 2000 contacts, can be programmed in *situ* and can download data remotely or through a mobile phone application, increasing their utility in remote areas or with species which are challenging to recapture.
4. We successfully trialled our loggers in a range of field realistic conditions, demonstrating that Bluetooth Low Energy is capable of logging associations in structurally complex habitats, and that changes in received signal strength can be equated to short range changes in distance between loggers. Furthermore, we tested the system on starlings (*Sturnidae vulgaris*).
5. The ability to include other sensors is retained in our prototypes, allowing for the potential integration of physiological and behavioural inference into social networks derived from our approach. Due to its open source nature, small size, flexibility of use and the active research currently being undertaken with Bluetooth Low Energy, we believe that our approach is a valuable addition to the biologging toolkit.

## 1. Introduction

Most animal social systems are heterogeneous; the extent to which animals will contact with each other will vary spatially and temporally [1] sometimes over relatively small time scales [2]. In order to accurately determine how population level social structure emerges from highly dynamic individual behaviour, it is essential to gather robust, accurate, high resolution empirical evidence of association behaviour [3]. However, systematic, disturbance free observation, particularly of highly mobile, nocturnal or small species, can be extremely challenging[3].

Understanding intra-specific associations is hugely important for understanding the processes underpinning survival, reproduction and disease transmission. How individuals associate with each other may mediate the flow of information transfer within a group [4] or establish social hierarchies[5], with multi-generational consequences[5]. The heterogeneous nature of social contacts also has consequences for understanding disease transmission, both within species of conservation concern[6, 7, 1] and hosts of zoonotic diseases[8, 9] or diseases of economic interest [10, 11]. Associations of interest may be inter- rather than intra-specific; animal associations are often embedded within a complex network of different species. Pairwise associations between two species may be modified by pathogen or predator mediated apparent competition [12], resulting in complex outcomes such as the apparent success of an inferior competitor in the presence or absence of a shared enemy[12]. However, it can also be challenging to assess the nature and frequency of these associations, with consequences for understanding demography and designing successful conservation programs[13]. A lack of a thorough understanding of these contact processes can even have serious, unintended consequences with a substantial conservation or economic impact [14]. Animals will also interact with their environment, often in conflict with human activities[15] or as a result of human behaviour, with potential consequences for survival or disease transmission[7, 8]. Proximity loggers, small devices worn by a target animal which log when another device is within a certain distance, can provide unparalleled insights into individual behaviour and associations. For example, such loggers have been used to identify inter-specific associations between cattle and badgers in relation to possible bovine tuberculosis transmission events [11], how contact patterns in raccoons relate to rabies transmission[16],to monitor whole herd movements for improved livestock management[17] or most recently to explore how sickness effects social encounters in wild vampire bats[18].

While the importance of accurately understanding contact behaviour is well established, limits have been imposed by technological capabilities. Classic approaches to determining contact between individuals often involve indirect approaches such as VHF transmitters[19], GPS loggers[20], and proximity loggers should provide a far more accurate picture[3]. Unlike methods which use spatial positioning to estimate associations, proximity loggers directly record the contact between two or more individuals. Proximity collars, one of the earliest examples of this technology (e.g. Sirtrack, New Zealand) record the length of time that loggers are less than a user defined distance apart from each other (e.g.40cm; [10]), thereby providing a duration of presumed contact[11], although this data is purely of a binary nature. While Sirtrack / Lotek proximity loggers have been instrumental in understanding the role of contact behaviour in a number of different systems[11], they are prohibitively large for many mammal species (collar weight is between 30 - 450g; [21]), and require the recovery of the collar in order to access the data. Despite these restrictions, such proximity loggers have been used successfully to determine the nature of contact behaviour between brushtail possums [10], white tailed deer [22] and raccoons [16], with consequences for understanding disease transmission.

As the majority of animals are smaller than the weight of a proximity collar, a key focus has been on creating smaller proximity loggers which will therefore be suitable for use on smaller animals. The first reduced size proximity logger was that of the Encounternet system. Originally 10g, modifications of these loggers achieved an impressive miniaturisation, with the smallest loggers weighing 1.3g[23], however this came with a significant reduction in battery life (only a number of hours) and are no longer available for use. Another recent approach to proximity logging in mammals involves the use of low frequency radio waves and a system of loggers connected to ground nodes[24]. The system is capable of efficiently and accurately monitoring associations between a number of individuals simultaneously, while also providing spatial information [24, 25, 18, 26]. In this system, mobile nodes are tracked by ground nodes providing spatial information, while encounter information is recorded by mobile nodes and stored until contact with a ground node [27]. While the encounter approach is comparable to the one we describe, ground nodes achieve contacts over a greater distance than BLE alone is capable of and long range movements can also be recorded. However, this system currently only works in Europe or America due to the frequency band used by the transceiver [26], is currently not available to researchers (pers comm Simon Ripperger), and the costs of implementing the system are unclear, as is the size and power consumption of the ground nodes. Therefore, while the BATS approach will answer questions concerning both proximity and long range movements of animals very effectively, inaccessibility remains an issue. Finally, a similar approach using a Bluetooth Low Energy (BLE) mesh has been described by [28] and by [17] where neighbour discovery using BLE is combined with LoRa technology to gather information on larger movements. Although the size of the proposed collar is not provided in either system, as a combination of BLE, LoRa and GPS collars are used, it is highly likely that these approaches have not been miniaturised for use on small (20g) animals, which has been the main focus of developing our system. Combining two radios (e.g. LoRa or NB-IoT) on the same chip would increase the weight while giving poor localisation accuracy (300m error; [29, 30]). LoRa and NB-IoT (two similar forms of long range low power wireless systems) have limited global coverage, particularly in Sub-saharan Africa where this system was originally designed to be used.

The majority of extant species are around 50g or less, limiting the proximity logger options that are available for use. In particular, species in the order Chiroptera or Rodentia are responsible for a large proportion of zoonotic diseases of interest, yet their average mass is 45g (3 - 491g; Pantheria dataset; https://ecologicaldata.org/wiki/pantheria). Current recommendations are that loggers do not exceed 5 % of the animals body weight for rodents and 8 % body weight for bats, although recent studies have exceeded this limit for short term studies [31]. Regardless, if loggers are too heavy, they will alter animal behaviour and provide inaccurate data. Therefore, our target was to miniaturise loggers capable of recording proximity data to 1g (not including housing or collar weights) in order to use on animals with a weight of 20g or more, staying within the restrictions of a 5 - 8 % of body mass weight limit. We do not include weights of housing (eg epoxy or sealant) or attachment method as these vary widely from species to species and will be subject to user experience with their study system.

Here, we present the system we have developed using Bluetooth Low Energy (BLE). Bluetooth Low Energy is highly efficient, capable of operating in high interference environments and is supported by modern phones and laptops, which means that user configuration does not require complex hardware[32], particularly in areas where infrastructure can be lacking. Our system consists of three components (Figure 1):

1. Contact loggers, which record the time stamp and the RSSI of the contact between individuals;
2. The gateways which store the logs downloaded from the contact loggers onto a microSD card;
3. A mobile phone application which allows real time programming, monitoring, and downloading of the loggers

**Figure 1.**
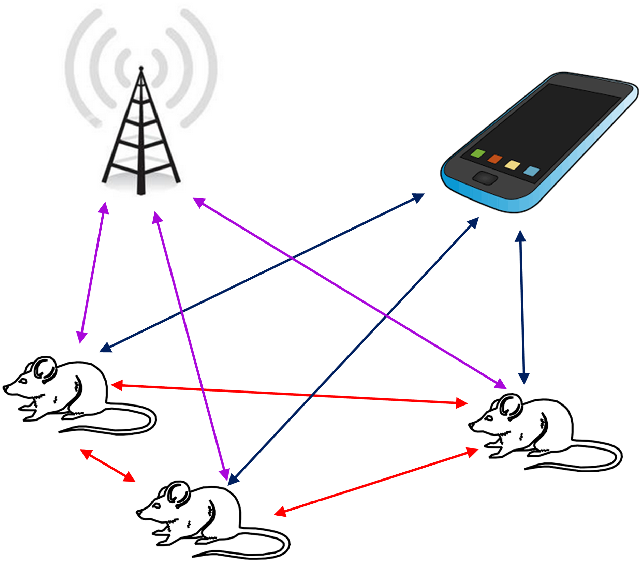
Schematic showing system set up. Coloured arrows indicate communication between different parts of the system. Loggers communicate directly between themselves when mounted on a focal animal (red arrows), downloading the stored data to either a gateway once a user determined threshold is reached (purple arrows) or a mobile phone application as and when a user desires (blue arrows).

First, we emphasise that the system is most appropriate for situations where users wish to investigate close contacts in a species where either the user can get close enough to download the data at some point (for example if the animal uses a nest) or gateways can be placed at strategic points for data download (e.g. known roost sites or within a closed grid), small scale spatial movements (up to 10m from a logger) are being monitored or when animals can be reliably recaptured to download data logs. Mesocosm studies would be particularly appropriate for this kind of system, although, as long as sufficient knowledge already exists about the species specific behaviour, open systems can also be used. This system will not provide information on animal movements over long distances, if animals spend a long time in a location where data download is not possible, or in situations where many animals are within a very small distances (<1m) of each other simultaneously; in these cases alternative approaches should be considered.

Our system is similar to previously described systems in some cases but also has some differences that we feel make it a valuable tool for answering specific questions. However, it should be noted that this system as is currently designed, will not provide the same long range spatial resolution as the BATS approach [26, 27, 24, 33], and indeed has not been designed to. Rather, our system concentrates on short range associations and spatial movements in an open source and easy to access system.

1. Our system is completely open source, low cost and readily available as we have concentrated on only using “off the shelf” components that can be easily accessed. By designing a mobile phone app for both Android and iOS devices, real time programming and monitoring of the loggers is easy and does not require specialist knowledge or equipment.
2. Contacts are directly stored on the chip of the logger, and downloaded once a user determined value is reached. This substantially increases the operational time of the logger by limiting contact between the loggers and the gateways, and allows the user to be circumspect about placement of the gateways which will download the stored data. How the user decides to place gateways or set download limits will depend on their knowledge of their study system; for example if the user believes that they are unlikely to regularly see their target animal, the download threshold can be set lower than in situations where the gateway is likely to regularly detect the mobile loggers.
3. Unlike Sirtek proximity collars, the IDs and the RSSI of the received identification are recorded, allowing fine grained differences in the association to be quantified and related to the potential nature and quality of the association;
4. The gateways have been designed to also operate under very low power meaning that they can be deployed in the field for months using relatively small batteries. This also ensures that they can be easily camouflaged in areas where interference or theft could occur and users can focus on just replacing loggers, or gateways can be placed in less accessible areas;
5. Similar to the BATS approach, the loggers and gateways are powered by rechargeable batteries, so loggers can be reused if recovered;
6. Loggers can be fully manipulated to match data requirements. Loggers can be set as ‘hidden’ where they do not broadcast their own ID but still scan for and record other IDs, ‘Advertise only’ where loggers broadcast their own ID but do not scan for other IDs, or ‘fully distributed’ where they both scan for IDs and advertise their own ID. Setting loggers as hidden stops stationary loggers from detecting each other, focusing data acquisition on the mobile loggers, while setting loggers as advertise only can substantially increase battery life. Loggers can be switched between these options in the field by using the mobile phone application.
7. The system has been designed to give complete flexibility to the user. Therefore limits can be set on the hours of operation (forced sleep during certain hours) and on which loggers are recognised by other loggers, again through use of the mobile phone app.

We recognise that some of the previously described systems have some but not all of the aforementioned points, however the open source nature, accessibility and low cost of this approach, we believe, makes it a valuable addition to the ecologist tool box.

## 2. Methods

### 2.1. General functionality of the system

Development of BLE (carried out by the Bluetooth Special Interest Group) is focused towards increasing energy efficiency [34]. BLE devices “advertise” their identification to their surroundings, the frequency of which is determined by the advertisement interval and a random back off interval which reduces potential collision risk between two loggers advertising at the same time[34]. Advertisements are also capable of holding some application data, meaning that the device is also able to publish data to its environment. Devices listen for advertisements by “scanning”, the length of which is determined by the scan window. The frequency at which a device scans is therefore its scan or measurement interval. The range over which BLE can transmit is determined by line of sight and the nature of any interference. Due to the miniaturisation of our loggers, transmission distances are considerably lower than those achieved by standard BLE (standard BLE can transmit up to 200m in open areas while our loggers transmit up to 10m as the board of the logger acts as part of the antenna for the signal and has been reduced below the optimum BLE operating distance). Complex habitat structure[11], particularly with a high water content, can substantially reduce the range over which the loggers can transmit[35]; therefore users need to consider the habitat within which their study species is moving, what constitutes a contact within their system before use, and ensure that loggers are calibrated. For example, when loggers are placed at floor level in thick undergrowth, transmission distances were reduced to 5m or less.

Our contact loggers scan and advertise within the default BLE schedule with user determined scan / advertisement parameters, storing any received IDs along with the RSSI and a time stamp. The loggers expose their unique identifier, amount of data logged and mode of operation in their advertisements, so other devices (eg. the gateway, mobile phones, tablets etc.) can access this data without connecting to the logger. Once the chip connects with a gateway it will download the stored data. If the connection with the gateway is lost before the full data transfer is completed, the data is not saved and will have to be downloaded again when the connection is restored. Once all the data is downloaded to the gateway, the contact logger memory is wiped. If the connection to the gateway is lost before all the data is downloaded, then the contact logger memory is not wiped and the data downloaded to the gateway is not stored. The data on the gateway is written to a microSD card which can then be retrieved by users at a convenient time. Data is written to the microSD card as a comma delimited file (.csv) for ease of onward processing. Contact data can also be directly downloaded from the loggers through the mobile phone application, as can programming the contact loggers to set the measurement interval, the mode of operation and the loggers unique identifier[32]. Loggers can be used in two different ways: As mobile nodes on moving animals which are restricted by weight, or as stationary nodes which are placed in the environment in a regular grid, do not have any weight restrictions and which provide spatial information, as well as inferring social contacts from proximity in space and time.

### 2.2. Contact loggers

The initial prototype was designed with a Silicon Labs BMG111 module[36], but subsequently we used BMG121[37] due to its smaller footprint[37]. The printed circuit board (0.3mm flex PCB) includes a detachable 6 pin TagConnect[38] connector which reduces the footprint required for programming and debugging, allowing us to maintain the small logger size. Battery terminals are at the bottom of the board. A voltage regulator is included to protect the chip from the high voltage from a fully charged battery. The PCB also contains a ground loop to tune the antenna and maximize transmit efficiency. A 47uF X5R decoupling capacitor is used to accommodate for sudden spikes in current the module needs for radio activity when smaller batteries struggle to provide sufficient current. In total, the chip weighs 230mg. Using the smallest prototype, the transmission distance is reduced compared to theoretical transmission distances as the length of the board is shorter [37]. The chips have two different modes to save on power, which can be combined. Chips can either operate at full power or low power which reduces both transmission range and battery consumption. The signal strength (TX power) on the BGM121 BLE chip is configurable, and as default is +8dBm. The low power option reduces this to −1dBm. Reducing transmission range can be beneficial in situations where users are only interested in close contacts, reducing the potential of contacting loggers further away, or increasing the time over which loggers are active is required. Loggers can also run as either scanning and advertising, where all loggers record the associations with other loggers, or as advertising only, where a network of stationary loggers can be used to infer contacts in space and time. After each advertisement, the module will listen for other devices to see if any other device wants to initiate a connection. If other devices in the area that are set to scanning want to connect, they wait for an advertisement, then immediately fire a connection request within that scan window. Using the advertising only setting can increase operation times dependent on the schedule being used. For example, using the smallest battery (10mA), when scanning and advertising on a 10 second scanning schedule (the highest time resolution we employ), battery life is 56 hours compared to 112 hours when advertising only. In comparison, logging on moderate accuracy (e.g. every minute) will require a current draw of 100uA, resulting in a runtime of 274 hours when advertising only, or 137 hours if scanning is enabled. Berkvens et al (2018) and Figure 2A-C demonstrates in more detail the relationship between battery life and measurement interval when full scanning is implemented, at full power. Most of the time the chip is in EM2 DeepSleep mode, where the timer continues to run but other parts of the chip are inactive. This can be supported by a 25mA battery for 333 days, and is 0.03% of the power consumption required for scanning [37]. By limiting the hours during which the logger is operational, the battery life can be extended (for example covering hours of activity for nocturnal animals, see Figure 2D).

**Figure 2.**
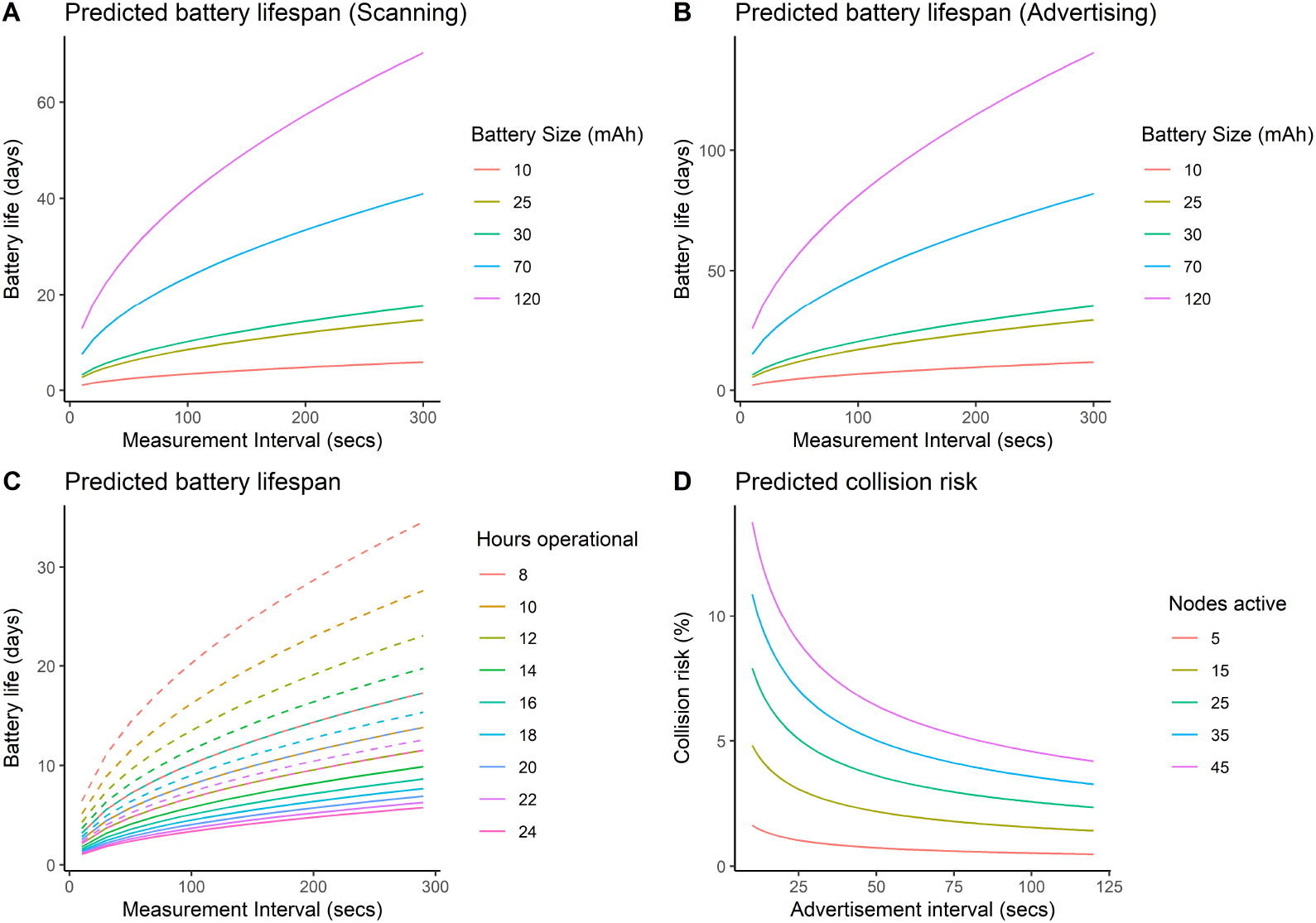
A: Predicted battery operating lifespan for a range of different battery sizes when scanning is enabled. B: Predicted battery operating lifespan for a range of different battery sizes when scanning is disabled. C: Predicted battery operating lifespan changes with hours operational. Loggers can be set to sleep for given periods which will increase battery lifespan. Solid lines show predicted lifespan when scanning is enabled, dashed lines show predicted lifespan when scanning is disabled. D: Predicted collision risk for given numbers of active nodes over a range of different scanning schedules.

In all cases we use a Lithium polymer (LiPo) battery to run both mobile and stationary nodes. Stationary nodes are a similar footprint to the mobile nodes, but are not restricted to 0.3mm boards (decreasing the cost of production) and include a battery connector for ease of use. The smallest batteries currently available are 10mA and weigh 0.4g. The choice of battery size for the logger will depend on the species being investigated and the time over which data is to be collected. Loggers can also be recharged and reused, extending the usability of a single logger.

### 2.3. Gateway

The prototype of the gateway is built upon the Nordic Development Kit for the nRF52840[39] which has full BLE5 support, an Adrafruit MicroSD card module is connected by soldering jump wires to the slot and inserting the wires in the appropriate connectors. A Adafruit GPS unit is also included to maintain an accurate time stamp. To facilitate power in the field, a 6600mAh Li-Po battery-pack is attached to a voltage-regulator module with its output wires soldered to the external power-input pins on the development kit. Alternatively the gateway can be powered with any rechargeable lithium battery with a micro-usb connector. The gateway will continuously scan for nearby loggers. When a logger is detected which holds data that exceeds the download-threshold, a connection is made and data is transferred to a temporary buffer. After the end of data is successfully detected, the gateway updates the clock on the logger to its own clock to ensure the timestamps on all loggers are synchronized. Both the nRF52 development board and the loggers have a 32kHz crystal at 20 ppm, with a drift rate of 2 seconds per day. The frequency at which the GPS unit updates the gateway is user determined, but an update every 4 - 8 hrs maintains 1 second accuracy across the whole system. Once the clock is successfully set, the gateway sends the erase-command which clears the data off the logger, after which the connection is closed and the data in the buffer gets written to the microSD card. When the connection is lost before the end of data is detected, the buffer gets cleared and no data is written to the microSD card. The data is also not wiped from the contact logger.

### 2.4. Mobile phone application

The mobile phone application is written in Dart using Flutter. The application uses the Bluetooth Low Energy capability supported on modern mobile phones to directly interface with the contact loggers. The app publishes a list of nearby contact loggers along with all the data that is in their advertisements (unique identifier, amount of data logged and operation mode). After a logger has been detected, a connection can be made which allows the user to edit parameters and to directly download the data. The binary data is automatically parsed into a CSV-file which can be opened by various spread-sheet apps present on the phone. The application can also emulate a gateway by automatically downloading and storing the data off nearby loggers, though this will put a severe strain on the battery of the phone and take longer; downloading through the gateway has been optimised for speed, with a download rate of 1.8 seconds per 100 logs. The unique ID for each logger is selected through the app, as is the scanning frequency.The app also allows users to select whether loggers are visible or hidden, whether they are advertising only, whether logs are limited to certain time periods and clears logs from loggers. The app also displays information such as the timestamp of the logger and the number of contacts stored on the logger. The app also displays the gateway when the gateway is functioning.

### 2.5. Trials

#### Battery life

We can estimate the average current draw of a logger by adding the charge consumed by all advertisements and the scan in a single cycle and dividing that by the length of the cycle. Sleep current has not been taken into account due to these currents being so small they become insignificant.

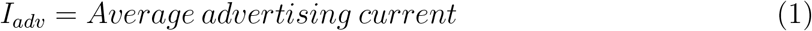

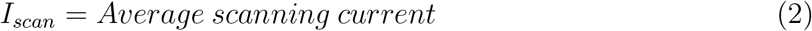

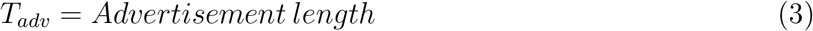

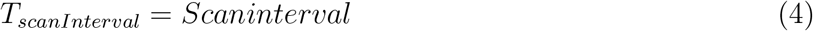

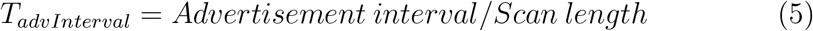

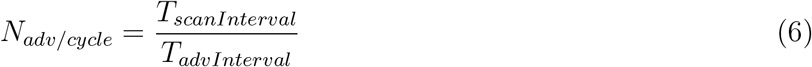

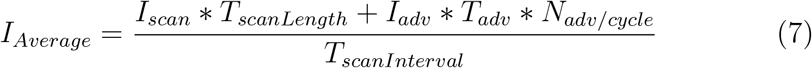

Multiple power-measurements were carried out in the Simplicity Energy Profiler to evaluate the accuracy of the model. Figure A6 shows the model applied on the BGM111 BLE-module, accompanied with actual measurements at set intervals.

#### Collision rates

Depending on the amount of Bluetooth Low Energy (BLE) devices in the immediate area, packet-collisions will occur. When two Bluetooth devices advertise simultaneously on a channel, both messages will render corrupt. This results in a chance that two or more loggers will not detect each other. The BLE-specification has measures in place to minimize these collisions but it is impossible to fully eliminate them. Following[40], we derived a model (eq A.1) to estimate collision rates depending on the advertisement interval, amount of nearby BLE-devices and the time it takes to completely transmit an advertisement (see Appendix A1 for details and Figure 2D for predicted collision risk for a range of nodes and advertisement intervals).

#### Contact logger tests

Initial tests were carried out to establish the range over which contact loggers could send and receive signals in a variety of different environments. First trials were carried out in Belgium to ensure that the tags were functioning as expected in open environments [32], with all following tests carried out at the field site in Morogoro, Tanzania during August 2018. We originally designed the loggers for use on *Mastomys natalensis*, a small rodent (~20 - 60g) that is widespread throughout sub-Saharan Africa. A prolific breeder[15], *M. natalensis* undergoes extreme population fluctuations in response to food availability and is a significant agricultural pest[15]. In addition,*M. natalensis* is the host for a range of zoonotic diseases including Lassa fever and plague[41], therefore understanding how social association behaviour influences disease transmission is of considerable interest for this species. Calibration tests were carried out in enclosed experimental mesocosms within which the preferred habitat of *M. natalensis* is maintained[41]. Tested habitats included thick grass (<30cm high) which had been cut and had all cuttings removed, thick grass had been cut, with cuttings left *in situ* and very long grass >2m; see supplementary data Figure A8 for images depicting the different habitats we trialled).

Initial logger calibration tests were carried out with both chips and batteries contained in plastic bags, and repeated after epoxy was applied to ensure that there was no negative consequences for the chip and battery from the epoxy. We found no evidence of epoxy application affecting the functioning of the Bluetooth chip, so continued all tests with loggers which had been coated in a thin layer of epoxy resin as deployment in the field will always require coating of some kind to ensure waterproofing of the loggers.

#### First validations

Two contact loggers were placed next to each other alongside tape measuring two metres. Loggers were each given a separate ID and the scan interval was set to 10 seconds. The gateway was set to reset the loggers after at least 10 contacts were recorded. Loggers were reset by the gateway to zero contacts, then the gateway was switched off. The contact loggers were left for 1 minute 30 seconds to record contacts. After recording contacts, the mobile phone application can be used to monitor the loggers and ensure that at least 10 contacts have been recorded by both loggers. The data was downloaded to a central .csv file stored on the mobile phone by selecting each logger in turn and downloading the data. The timestamp at which the data was downloaded is recorded in the data file. One contact logger was then moved ten centimetres along the measuring tape, the gateway was turned on to reset the loggers and the process was repeated. Each time the data is downloaded from the logger it is appended to a single .csv file for ease of management, as well as creating separate logger specific download files. The data was also downloaded to the gateway each time the loggers were reset but for ease of handling we advise using the mobile phone application as the contact loggers can be monitored in real time. This process of moving one logger was repeated every 10cm for one metre, after which we moved the logger every 20cm for the next metre. This process was repeated for loggers without epoxy, loggers with epoxy and loggers mounted on laboratory gloves filled with 48ml of water to mimic one of our focal animals (see Figure 4 and 5 for declines in RSSI over distance for each trial).

**Figure 3.**
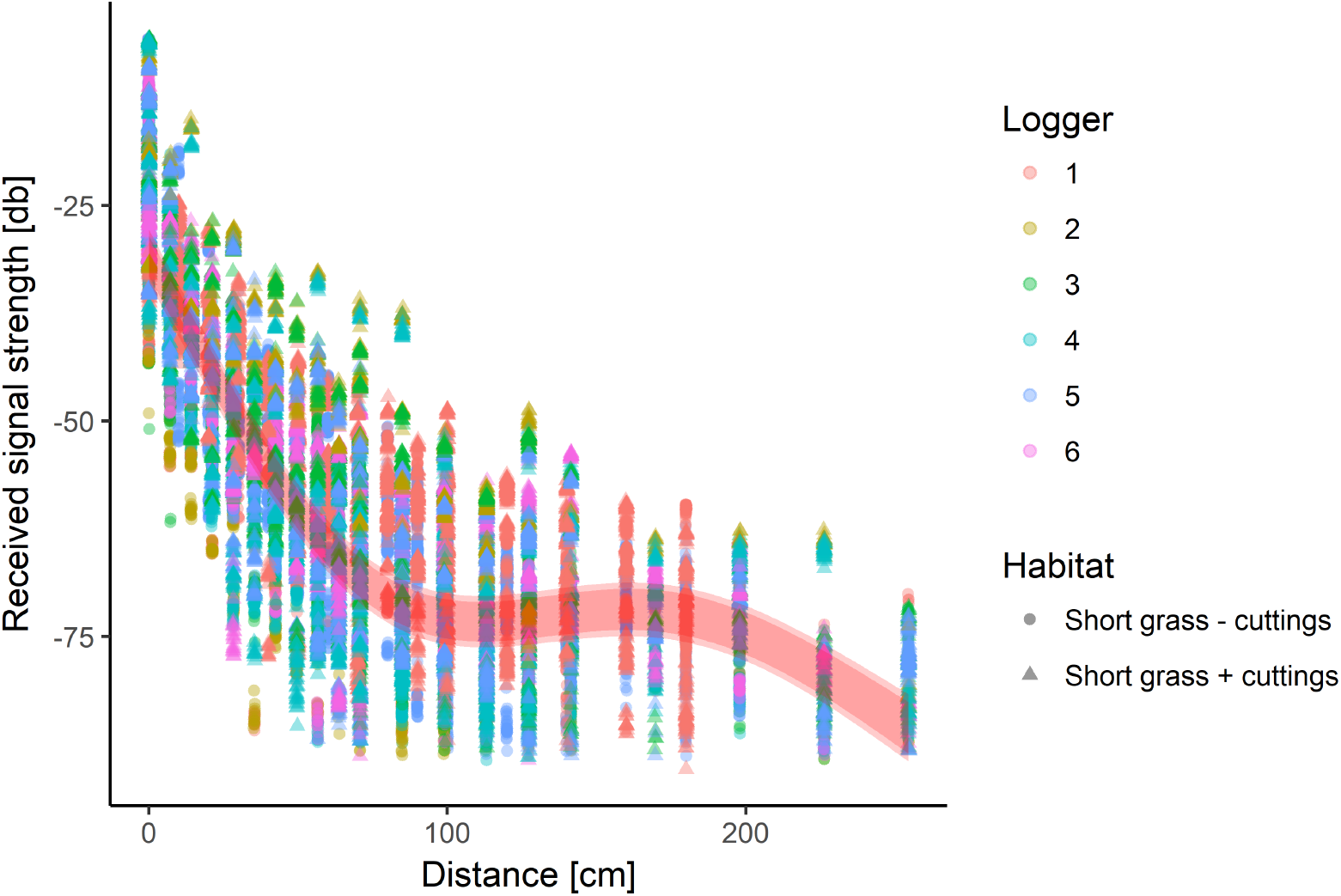
Relationship between RSSI and distance for two habitat types and 5 loggers (all with epoxy applied). Points indicate raw measurements in different habitats, ribbons indicate predicted relationship between RSSI and distance returned from the model, pale ribbon indicates the 95% simultaneous confidence intervals.

**Figure 4.**
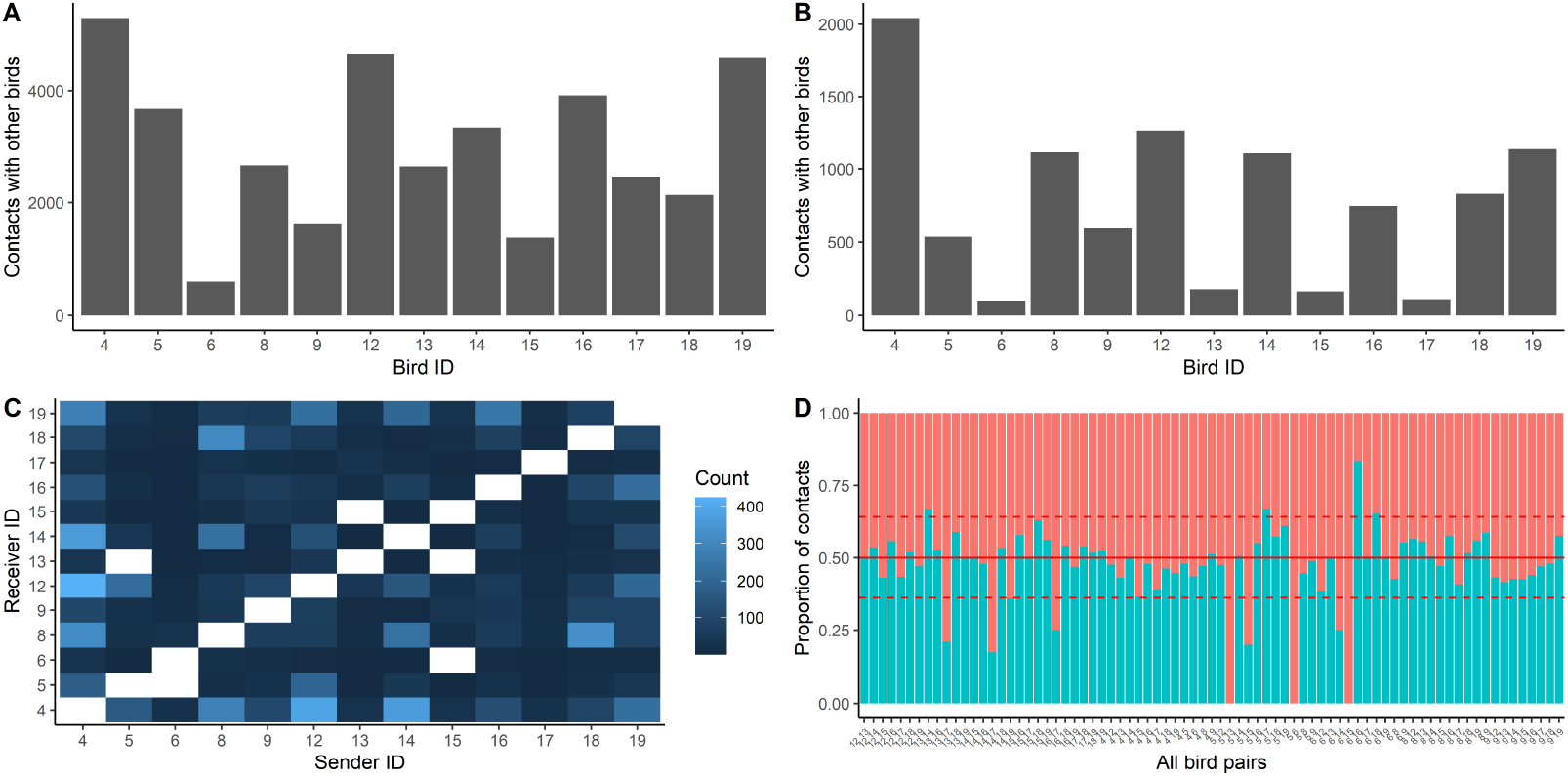
Contacts recorded with other birds for each bird for A: Contacts (RSSI ¿ −60) where birds are within half a metre of eachother and B: Close contacts (RSSI ¿ −50) where birds are within a few centimetres of each other. C: Plot showing registers on each pair of loggers, fill shows the count of logs recorded by each logger in the pair; D: Proportion of total contacts recorded by each logger in a pair. Solid red line indicates 0.5 where both loggers have recorded equal logs of eachother, dashed lines represent the standard deviation. Logger pairings which fall outside of the standard deviation indicate where one logger in the pair recorded more / less logs than the other logger.

**Figure 5.**
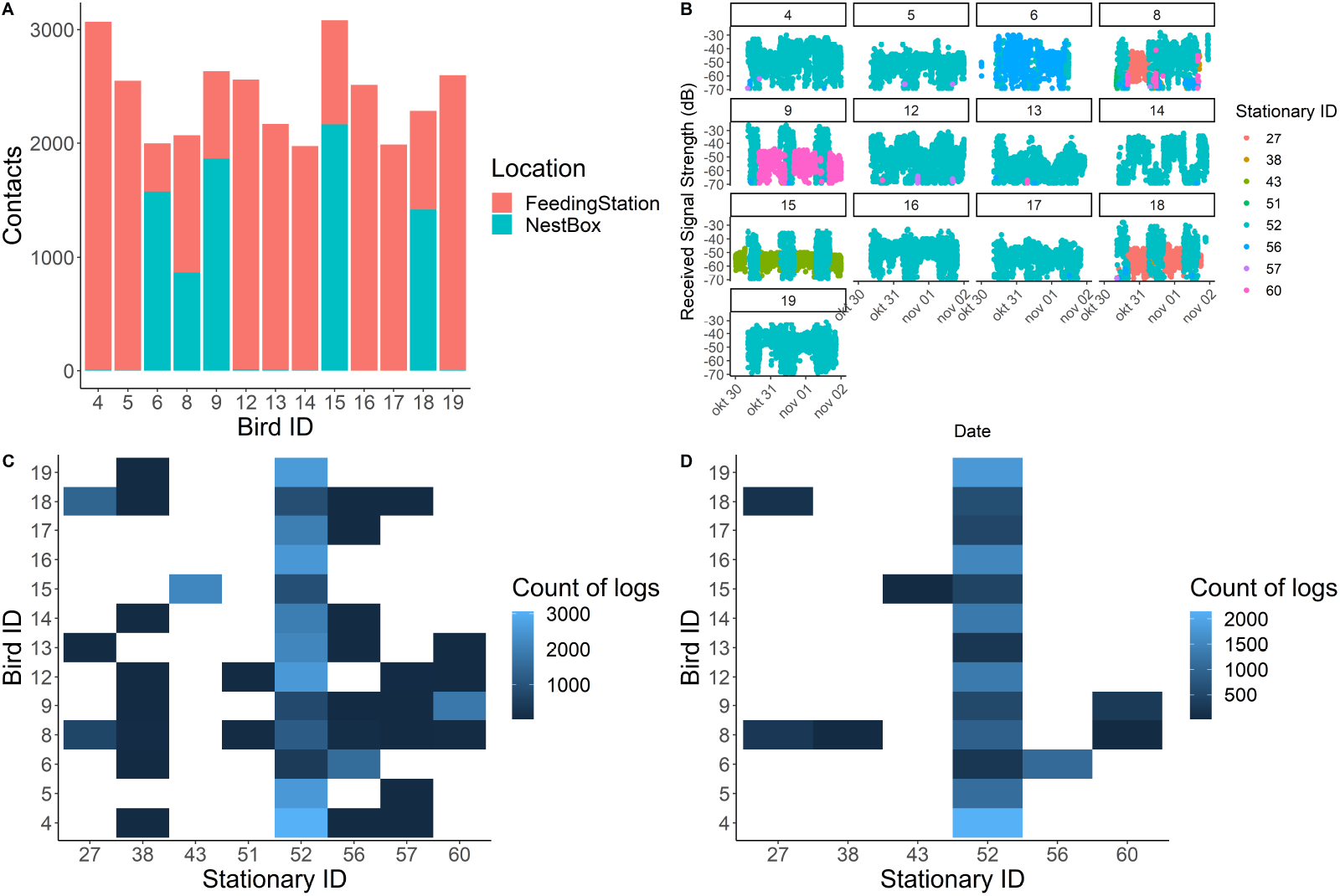
A: Number of logs recorded on stationary loggers for each bird separated by whether the stationary logger was placed at the feeding station or a nestbox. B: Temporal fluctuations in contacts between stationary and mobile loggers during the course of the experiment. Logs are filtered to only consider contacts with an RSSI of −70dB or more. C: Heatmap showing logs by nest box and feeders for each bird when considering contacts (RSSI ¿ - 70dB). D: Heatmap showing very close (¿-50dB) contacts between birds and stationary loggers.

#### Grid validations

Rutz et al. 2015 describe a detailed approach to calibrating animal borne proximity sensors which combines a thorough documentation of the distance signal strength relationship across the three-dimensional environment the focus animal will move through[42, 24] with statistical models and computer simulations[42]. Furthermore, the size and behaviour of the tagged animal will also influence the relationship between signal strength and distance. Loggers attached to arboreal mammals will detect each other over increased distances when ascending a tree compared to when moving terrestrially in long grass, and the water content of the animal itself may also influence the range of BLE transmission[35]. Due to these considerations, accurate calibration, tailored to the specifics of both the focal species and the habitat in which the focal species move is vital. We designed a calibration routine which was suitable for our specific habitat (see supplementary data A3 for a detailed description), allows the simultaneous testing of five loggers, and would be appropriate for any terrestrial, non-arboreal species. Two measuring tapes are laid out in a cross, with distances marked on them as described in Supplementary data A3. One logger is placed at the centre of the cross and remains there for the duration of the test, while the four other loggers are placed on each arm of the cross. As each logger is moved along the arm of the cross, it will move a set distance from the other four loggers (see Supplementary data A3). The same protocol is used as described above; loggers are set to advertise every 10 seconds, and loggers are moved after one minute 30 seconds again. The data is downloaded from all five loggers to the mobile phone application between each movement. This was repeated twice in two different representative habitats in our study area (thick grass without cuttings and thick grass with cuttings).

### 2.6. Statistical analysis

Theoretical models for battery life and collision rate were carried out in Matlab. All statistical analysis was carried out in R (R core development team, version: 3.4.4). The relationship between RSSI and distance was validated using a linear model (mounted logger trials), as was the relationship between distance moved and average contacts recorded. The relationship between RSSI and distance for the grid validation was modeled using an additive model with a gaussian distribution, including a smoothed term for distance and habitat type and logger ID as fixed effects. Residuals were checked visually for normality.

### 2.7. Field realistic trial

Loggers were initially tested in a field realistic trial on a captive colony of common starlings *Sturnus vulgaris* enclosed in a large aviary (50m x 10m). We chose to test the loggers inside an aviary as that way we would identify periods with missing logs as a consequence of system malfunction rather than missing animals, and the field test was carried out on starlings based on availability. We tagged 15 birds (8 males and 7 females) with Proxlogs attached as backpacks sealed in epoxy resin. Loggers were all below 5% of the birds’ body weight. Eight nest boxes were placed in the aviary, with loggers placed underneath the box and a Bushnell wildlife camera placed in front of each box. Cameras were set to record for 30 seconds after being triggered, allowing contacts between birds and stationary loggers placed at nest boxes to be verified. It is not possible to observe the birds directly as the presence of an observer is too disturbing for the birds, and this way we were able to observe for 24 hours a day. Birds were provided with clean water for bathing and a feeding station. The feeding station also had a stationary logger. Mobile loggers on the birds were set to ‘scan’ every 120 seconds for the full 24 hour period. Stationary loggers were set to ‘hidden’ so they were able to record contacts with the mobile loggers but did not record other stationary loggers. The scan schedule for the stationary loggers was set at 120 seconds for the full 24 hour period. Data was downloaded automatically through the gateway which was placed outside the aviary in the centre. This is adjacent to the feeding station so likely to detect all birds regularly, but was able to download from all stationary loggers in this position. The birds were checked every day for signs of problems, after 3 full days of logging birds were recaptured, loggers removed and birds checked for any sign of injury.

### 2.8. Acquiring the loggers

Users interested in discussing whether the loggers are appropriate for their study system or question can contact the authors on for more information on accessing and using the system.

## 3. Results

### 3.1. Tag functionality

#### Battery life and collision rates

The choice of battery size is constrained by the size of the focal animal. With the smallest batteries (10mAh, 0.4g), and a measurement interval of 10 seconds, we predict a battery life of 2.3 days. This can be extended by either increasing the measurement interval (e.g a measurement interval of five minutes will extend battery life to 12.8 days) or by only logging associations during the period of known activity (figure 2C), which will increase the predicted lifespan. Our theoretical predictions of battery life similar to those we experienced in the field during our trials and matched actual measurements (see Appendix 1 Figure A6; [32]).

Figure 2D plots expected collision rates based on the model from [40]. We observe an elevated amount of collisions when enforcing a low scan-interval and a large number of nodes (high data-resolution). This is expected as more advertisement-transmissions are required when scanning frequently, thus resulting in a higher congestion of the air-space. This extreme example highlights that collision risk will be higher if you are expecting a large number of animals (more than 30) to be within a few metres of each other and you have a high scanning rate. In these situations we would suggest that another system may be more appropriate.

#### Logger function in field realistic conditions

Loggers were tested in field realistic settings to determine whether using Bluetooth Low Energy would be suitable for animal borne proximity loggers. Encouragingly, we found that our system was able to detect advertisements in a range of habitats representative of our focal species’ preferred habitats. The range over which we were able to detect contacts differed with habitat type and between loggers (Table 1), reinforcing the importance of calibration for effective logger use.

#### Validations

The relationship between distance and received signal strength (RSSI) is variable depending on both the logger itself and the habitat within which the logger is moving. Adding the loggers to gloves filled with water to mimic a rodent body did not cause any change to signal transmission in the three different habitats (Figure 4). Signal transmission declined more steeply with distance in the very long grass than in either the cut grass with cuttings removed or the cut grass with cuttings retained (F_2468_ = 39140; short grass no cuttings: −44.7 ± 0.4; short grass + cuttings: −43.0 0.4; uncut grass: −50.1 ± 0.3, adjusted R2 = 0.98; Figure 5).

#### Grid validation

The additive model accounted for 73 % of the variation in received signal strength. We found significant variation between loggers (Table 1), for example, logger 4 consistently recorded lower RSSI values than other loggers. Distances below 30 cm, which could constitute a “contact” in our system, were predicted by RSSI values of an average of −27 (95 CI −10.8 - −43.6) dB (Figure 5). However, we did find occasions where dyads of associations were not registered (i.e. contacts were recorded on one logger but not the other logger). The extent to which this occurred increased with distance (F_478_ = 25.8, change in position: −0.03 ± 0.007); at the shortest distance loggers had an average of 3.5 (2.3 - 4.0 95 CI) contacts compared to 3.0 (1.5 - 4.0 95 CI) as distance increased.

#### Gateway

Increasing the height of the gateway increased the distances at which the gateway was able to connect with the loggers. If the gateway was moved from 15cm off the ground to 1m off the ground, the distance at which it could receive loggers increased from 5.5m to 11.7m. Raising it a further metre from the ground increased the distance to 18.2m due to improved line of sight. It is therefore advised to consider the distance over which tag download is required when placing gateways. Signal strength at the gateways can be increased by the addition of an antenna, increasing the potential coverage of the gateway. However, this is beyond the scope of what is currently developed for the system, and has not yet been tested.

### 3.2. Field trial results

#### Logger success rate

Starling experiment: Of 15 birds fitted with a logger, 13 retained the logger in working order for the full 3 days of the experiment while two loggers failed. As loggers were sealed with epoxy, logger recovery is not possible and therefore it is not known why loggers failed.

Mobile loggers mounted on birds recorded a total of 103029 contacts, of which 39132 (37%) signify actual contacts (RSSI greater than −80dB), and 9966 (10 % of the total logs and 25% of the contact logs) would be considered close contacts (RSSI greater than −50dB implying that the two loggers are very close to eachother). Contacts were fairly evenly distributed between birds (Figure 4A) but when concentrating on close contacts it is clear that some birds had a lot more close contacts than others (Figure 4B). Contacts should be logged twice, by both individuals involved in the association. When considering close contacts, in the majority of cases logs on each member of a pair match each other (Figure 4C), although there are some cases where one bird logged contacts and another didn’t. For example, bird 5 did not log any contacts with bird 6, but bird 6 did log one contact with bird 5. Birds logged eachother more similarly when considering individuals with more logs (Figure 4D); bird pairings with very uneven logs were all those which had a very small number of logs (less than 10 logs in total between both birds).

Stationary loggers recorded a total of 31,629 logs, of which 14,368 could be considered very close contacts and would indicate the bird is on the feeder or in the nest box. The vast majority of logs were recorded by the logger placed at the feeding station (23,602 logs, 12,587 close contact logs; Figure 5A). Loggers were placed at the 8 nest boxes distributed in the aviary; while all nest box loggers recorded some associations, not all boxes recorded close associations suggesting that birds did not use all boxes (Figure 5A). Associations with the feeder and nest boxes varies between birds, with some birds staying in the vicinity of the feeding station at all times, while others split their time between the feeding station and nest boxes (Figure 5B, 5C). Some nest boxes were also more popular than others, with boxes 57 and 51 appearing to have few contacts and no close contacts (Figure 5C, 5D). Combining stationary and mobile logger data revealed that birds were sharing nest boxes overnight (e.g birds 8 and 18).

Comparison with camera footage: Comparing logs with the camera trap footage revealed that logs of RSSI −60dB and greater corresponded to a bird interacting with the box (sitting on the perch, being inside the box, or sitting inside the box looking out). An RSSI of −50 dB or greater corresponded with a bird being inside the box. Nestbox loggers recorded 7094 associations at −60 dB or greater, of which 6 (0.08%) could not be matched with either direct camera trap footage of a bird entering or leaving a box, or were logs of the periods between which birds were seen entering or leaving a nest box. Camera trap footage can be associated directly with 46 (0.7%) of logs, indicating birds either sitting on the perch outside the box or entering and leaving the nest box. In some cases the camera was clearly triggered by a bird entering or leaving the box, but the bird was either not visible (but box shaking and a close contact log recorded) or was just visible. In 30 cases (0.42%), logs show associations with the boxes that are not detected at all by the camera trap. It was rare that camera trap picture quality was sufficient to ID a bird, but the ID of the bird could be determined by cross referencing the logger ID with the camera trap footage. Logger data gave additional data that would not be possible to retrieve from camera traps alone. The camera traps often missed an entry or an exit, or the ID of the bird was not visible so the duration and ID of any birds association with the nest box would be unknown; 80% of the logs between nest boxes and birds occurred over night or when a camera had missed a bird enter or leave. After removing camera trap footage involving a bird with a broken logger, there were 8 occasions (0.01% of associations) where birds were caught by the camera trap at a box without any corresponding logger data. While camera traps do record interactions and behaviour that would not be inferred from logger data (for example antagonistic interactions between two birds at a nest box), loggers also captured behaviour that was missed by cameras, such as birds sharing a box when the entry of one bird was not captured on the camera traps. Furthermore, the loggers provide reliable information about the ID of the animal involved in the associations which was not always possible to determine from camera trap data, and would not be possible from observations given that the most interesting associations were occurring at dusk and dawn.

Comparison of the stationary, mobile and camera trap data shows that the majority of bird associations took place away from nest boxes. Of nearly 10,000 close contact logs that were recorded, 40% were in close proximity of the feeder, 0.8% were in close proximity with nest boxes and the other 59% were elsewhere in the aviary.

## 4. Discussion

Common analytical tools used to explore animal contact networks, such as graph theory, are known to be highly sensitive to the sampling effort carried out to define the network [2]. Missing associations can have significant consequences for some topographical statistics [43], therefore accurately quantifying associations is vital for parameterizing many network analysis approaches[43]. Furthermore, the ability to record the behaviour of the most species rich body weight classes in birds and mammals depends on either battery miniaturisation or reduced energy consumption of such tags[21]. Here we present a novel approach to determine contacts between wild animals using extensive miniaturisation and Bluetooth Low Energy, a form of wireless communication which is currently under active development. To date, weighted automated social network data on small animals derived from proximity loggers are sparse due to the size constraints imposed by the loggers themselves[23]. While approaches using RFID readers have become more popular in recent years, these can only record associations within the presence of a reader, which may involve altering animal behaviour to record the association (for example providing feeders or nest boxes to record associations). While these experiments can reveal fascinating insights into animal behaviour, our live experiment showed that the majority of associations actually took place away from the feeder or a nest box, showing the utility of proximity detection systems for providing a continuous log of animal association behaviour [18].

We experimentally tested our system performance by tagging 15 *Sturnus vulgaris* in a large aviary, with stationary loggers placed at a feeding station and eight nest boxes. Two loggers failed shortly after attachment but the others collected data for the full period of the experiment. Coverage was very consistent throughout the experiment, with data collected at a high temporal resolution. We found little evidence of substantial data loss due to collisions, with most logs mirrored on both loggers particularly when considering close contacts only. Logs deviated from being very similar when very few logs were recorded, which may suggest that these associations were only fleeting rather than data loss due to collision risk. Comparing the camera trap and logger data showed a high accordance between the two. Logger data and camera trap data was able to be matched 99% of the time, although both forms of surveillance provided different forms of additional data. While bird interactions with each other and the nest boxes were observed on the camera traps, the bird ID was often hard to identify from rings due to the picture quality or the time of the photo (after dark and therefore not in colour), and had to be inferred from the loggers. Furthermore, camera traps missed key moments like birds swiftly entering and leaving the boxes while loggers provided a consistent record of bird presence in the boxes and with each other. In contrast a very small number of associations were recorded at nest boxes that were missed by the loggers (less than 1% of the total associations). In this experiment the loggers were fitted with 10mAh batteries, which were predicted to operate for 88 hours (given a 24hr runtime period) and were still operational at the end of the experiment (64 hours). Our approach includes a range of battery options which will allow the development of loggers with a minimum weight of <1g depending on mounting options. However, while such small loggers increase both the species and individuals within species in which proximity behaviour can be explored, it should be noted that, as with all these systems, we are still only able to monitor a subset of the population due to trapping biases and individuals which do not meet the minimum weight requirements[43], and when monitoring such small animals, powered systems will always have some limitation to runtime. By incorporating different energy management regimes, we have increased the potential runtimes that can be achieved, increasing the data which can be gathered. Nevertheless, miniature proximity sensors, produced with off-the-shelf components such as we have used here provide an inexpensive and lightweight approach to monitoring association behaviour between wild animals.

As currently described, our system would be most appropriate for monitoring proximity behaviour of species which are either enclosed within a space that can be easily monitored with gateways (e.g. mesocosm experiments), or regularly pass or return to known points in order to download contact data. Our chips were able to consistently communicate with each other, the gateway and the mobile phone application in the field, validating the approach. The addition of epoxy did not change the effectiveness of the approach, suggesting that sealing to ensure that loggers are safe from damage will not adversely affect the system. As the antenna is contained within the PCB, we do not expect to see changes in RSSI in response to antenna manipulation as has been described from other systems[44, 24].

We predict a longer battery life than that described for the Encounternet system[23], with a similar temporal resolution. However, our system has additional flexibility built in that can be used to extend the battery life, such as by setting the logger to sleep during certain periods of known inactivity, or by deactivating scanning. Due to our configuration of data storage on the chip, our loggers are able to store up to 2000 contacts before becoming full, increasing the period during which focal animals can be away from the gateways before data loss occurs[25] and allowing for contacts to be recorded when animals are in unknown locations. Alternatively, in systems where a relatively low number of encounters are expected, gateways can be set to download data from loggers when a lower number of contacts are stored, reducing the risk of losing data due to tag malfunction or loss of a focal animal[44]. Collision risk increases when a large number of loggers are in close proximity to each other; in systems where this is likely it may be preferable to increase the measurement interval to reduce the likelihood of collisions. While we recognise that this is not ideal, using the loggers within an enclosed aviary with 13 birds and 9 stationary loggers still resulted in high resolution social and spatial information. We have also made the system as simple to use as possible, the mobile phone application makes it straightforward to monitor and adjust detection settings in real time, including after loggers have been attached to focal animals while data is directly downloaded as a .csv file which can easily be manipulated for analysis.

The greatest challenge with analyzing proximity data is the conversion from RSSI to distances between animals[42, 24]. When moved at ground level in structurally complex habitats, loggers were able to detect the presence of other loggers over a range of distances below one metre. Between two to three metres, this relationship was less clear and differentiating distances was no longer possible, although loggers were still able to make contact. This is lower than distances reported from the “Encounternet” system [23] and [24], but our tests only considered movement for terrestrial rather than aerial species in structurally complex habitats. In our live tests, comparison of logging data and camera trap data revealed that RSSI’s of −50dB or greater were consistently aligned with very close associations (within a few centimetres) in line with our tests and could therefore confidently be assigned to a contact. For larger animals the RSSI at which a contact is assigned may be lower and would require calibration. Detection distances between loggers will increase substantially in open space, if animals move vertically as well as horizontally, or with the addition of an antenna, which may be appropriate for other study systems. In our case, as we are interested in characterising associations involved in direct virus transmission, contacts over short distances are desirable. However,the open source nature of our approach means that other research groups can test the extent to which antenna will increase transmission distances if appropriate for other case uses. In our live test we also found that the gateway was able to reliably download loggers that were up to 30m distant, reflecting the increase in transmission distance when both loggers and the gateway are placed at least 1m from the floor. In situations where users may want a larger gateway coverage, additional gateways can be used to download data and will not interfere with eachother. Although our system showed fairly stable declines in RSSI over short (<1m) distance within different habitat types, there was considerable variation between the different loggers and this needs to be accounted for in the calibration. We present an approach, derived from [42], which allows users to easily and relatively swiftly calibrate a number of loggers at one time; this is essential to estimate distance categories which reflect reality in the focal system[24, 42, 4].

The miniaturisation of biologgers is an exciting development for researchers who want to understand how association behaviour influences a range of different processes. How animals interact with each other is fundamental to understanding both the biology and behaviour of animals[3], with consequences for disease transmission[6], gene flow[2], information transfer[4] and resource exploitation [45]. The utility of proximity loggers is not restricted to mammalian or avian species; with sufficient miniaturisation loggers can also be applied to large invertebrate species, and current logger sizes would not preclude the use of these loggers on many reptilian species. Automated processes with remote access availability will increase the range of species such information can be collected on as the majority of vertebrate species are either small, cryptic or impossible to observe directly in the field[21, 13].

### 4.1. Future directions

In recent years tracking technology has passed important thresholds in both the size of the logger and the resolution of the data being collected [21]; miniaturised proximity loggers will not only allow an increased quantity and quality of data to be collected, but also allow the addition of other sensors to augment the proximity data being collected[46], providing an integrated view of the animal and its environment[46, 21]. For example, [25] demonstrated how including an accelerometer provides insight into the behaviour of tagged bats during monitoring with proximity sensors as well as incorporating an elegant way of restricting energy use to periods of activity. Temperature loggers may be useful to indicate arousal from torpor, or to equate association behaviour to environmental conditions[46]. The addition of other sensors to our loggers is easily achieved, although the energy requirements and additional weight of any sensors needs to be taken into account. Gateway development is currently underway to include the ability to download data remotely by accessing mobile data networks as well as creating a meshed “network” of gateways, extending the range over which loggers can be reliably downloaded. This approach would allow gateways to communicate between each other, offloading data to a single “master’ gateway. Finally, recent improvements in range for BLE transmission means that data may now be collected over a larger spatial area or for a greater range of research questions. Our approach complements the similar approaches designed by[24, 26, 28, 17], and adds another method to the growing toolbox of biologging approaches, particularly because the open source, low cost nature of our approach means accessing our system should be more achievable for a range of different users.

## 5. Acknowledgements

We thank Christine Talmage, Geoff Sabuni and Shabani Lutea for their help during fieldwork in Tanzania, and Dragan Subotic and Kwinten Schram for help during system development. This research was funded with a Global Minds Small Research Grant (GMKP0101). LK is a Junior Fellow with the Fonds-Wetenschappelijk-Onderzoek (FWO Research Foundation) which has co-financed this work (Grant nos: 1220820N and 1513519N). We thank Marcel Eens, Geert Stevens, Peter Scheys and Frank Adriaensen for assistance with carrying out the starling experiment.

## 6. Data accessibility

Logging data and the R code used to download and manipulate the data will be made available as example data for R functions / R package to manipulate logger data.

## 7. Author contributions

LK, HL, RB, WM and IHO conceived the study design. LK, RB and IHO designed the methodology, LK, IHO, AM and CS participated in the fieldwork, LK and IHO analysed the data, LK and IHO led the writing of the manuscript. All authors contributed critically to the drafts and gave final approval for publication. LK and IHO contributed equally to the writing of this manuscript.

## Appendix A. Supplementary data

## Appendix A.1. Collision rates

Statistical model derived from [40]

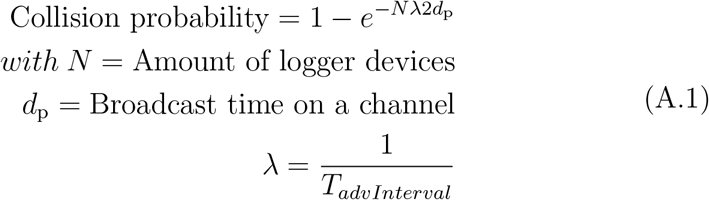

## Appendix A.2. Power measurement validations

## Appendix A.3. Calibration grid for validation

The calibration was carried out by laying out a grid as shown above. Loggers were placed in each of the five place marked positions. Loggers were moved along each arm of the cross with the central logger staying stationary. Loggers with distance A separating them increased by 10 cm each movement until they were one metre apart, at which p√oint they increased by 20 cm each movement. Distance B was calculated as 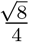 * A, distance C was calculated as 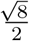 * A. In order to maintain these distances loggers were moved 7.07 cm along each arm. All calculations were carried out in R (R Core Development Team, v 3.4.4)

## Appendix A.4. Images of the field site and loggers

## Appendix A.5. Set up of field experiment and results

**Table 1:** Estimates and standard error from additive model including distance as a smoothed term. Distances were calculated using the grid calibration validation.

| **Model** | **Estimate** | **Standard error** |
| --- | --- | --- |
| Uncut grass + cuttings | -54.5 | 0.12 |
| Uncut grass - cuttings | -4.9 | 0.10 |
| Logger 1 | -2.6 | 0.16 |
| Logger 2 | -0.7 | 0.16 |
| Logger 3 | -4.5 | 0.16 |
| Logger 4 | -0.3 | 0.16 |
| Logger 5 | -1.3 | 0.16 |

**Figure A.6.**
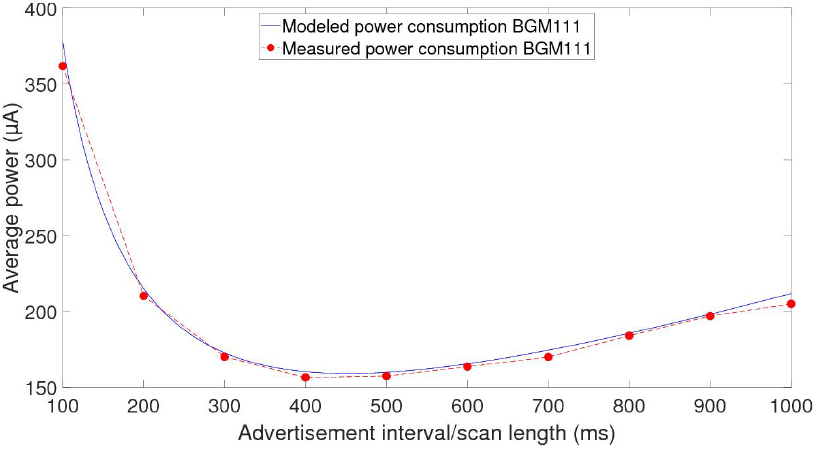
Comparing modelled power consumption with actual power draw as advertisement interval increases. At all times actual power draw closely approximated that predicted by our model

**Figure A.7.**
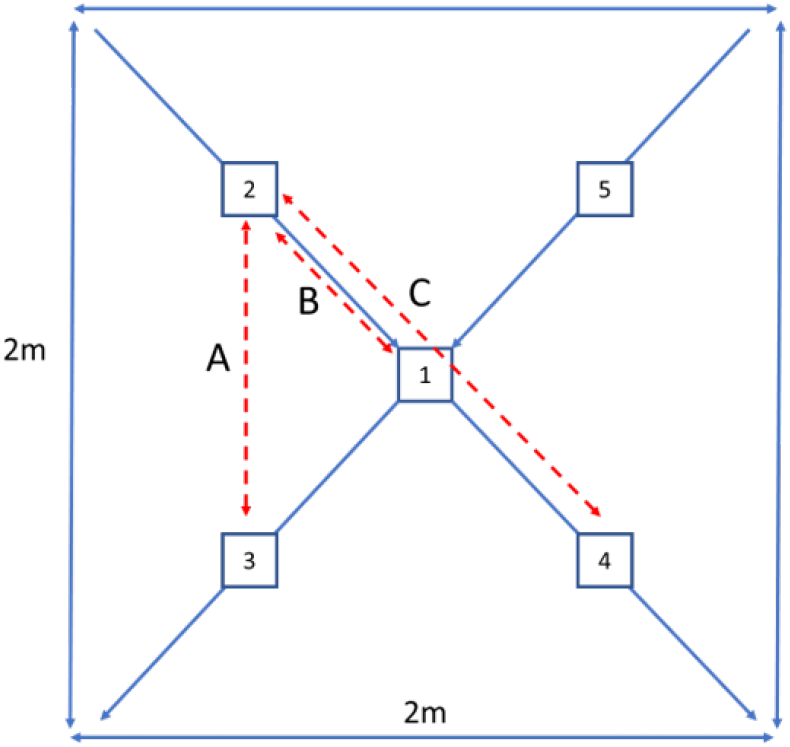
Grid layout for performing calibrations

**Figure A.8.**
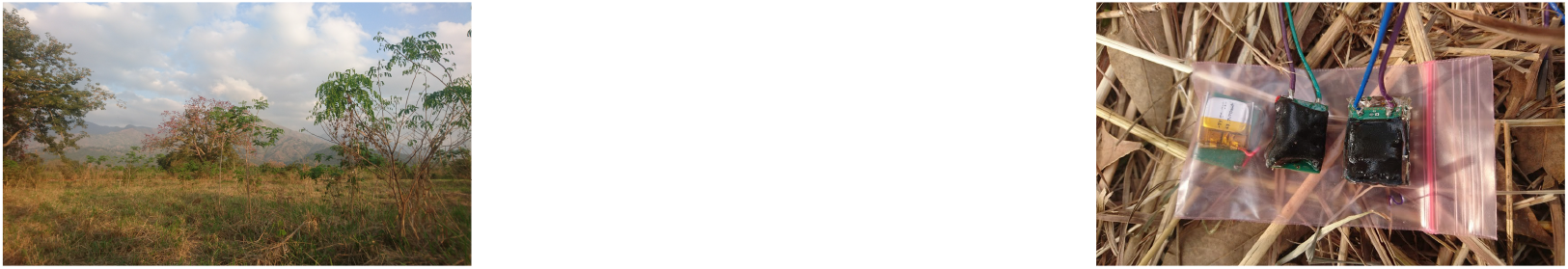
Top: Landscape showing habitat in which trials were carried out; Bottom: Prototype loggers (two with epoxy) used in trials

**Figure A.9.**
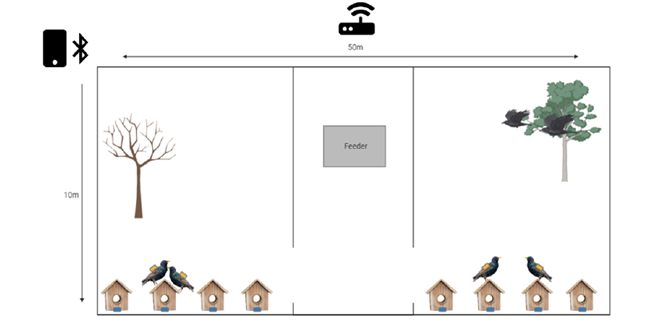
Layout of the starling aviary for the field trial. The feeding station is in the centre of the aviary, with a stationary logger attached. Nest boxes are arranged along one side of the aviary, with a camera trap placed in front of each. Blue squares represent the stationary loggers while yellow squares represent the mobile loggers. A gateway was placed adjacent to the feeding station but on the outside of the aviary to allow easy access and ensure that mobile loggers would be downloaded. In addition, data could be read and logger settings adjusted through the use of the mobile application.

**Figure A.10.**
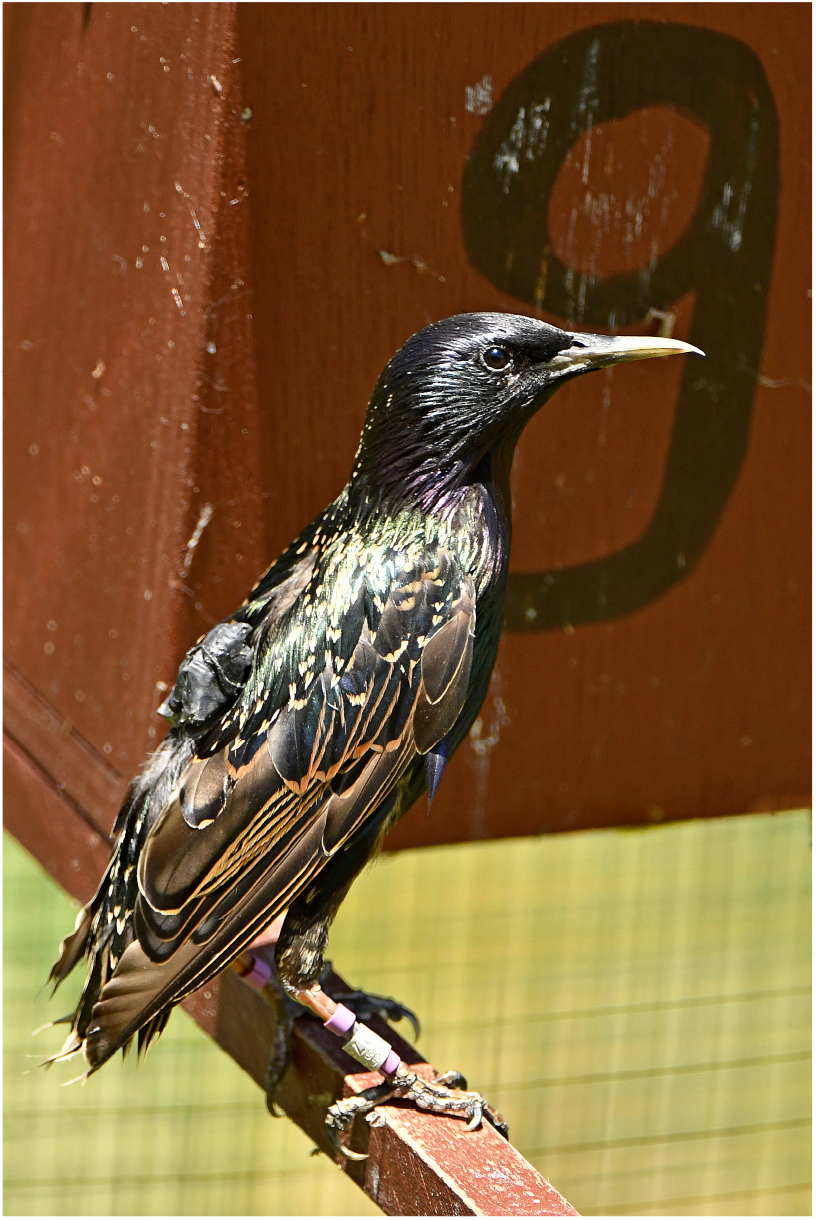
Picture of starling with a logger attached as a backpack.

## References

[1] K. L. VanderWaal, H. Wang, B. McCowan, H. Fushing, L. A. Isbell, Multilevel social organization and space use in reticulated giraffe (Giraffa camelopardalis), Behavioral Ecology 25 (2014) 17–26.

[2] C. Tentelier, J. C. Aymes, B. Spitz, J. Rives, Using proximity loggers to describe the sexual network of a freshwater fish, Environmental Biology of Fishes 99 (2016) 621–631.

[3] J. Krause, S. Krause, R. Arlinghaus, I. Psorakis, S. Roberts, C. Rutz, Reality mining of animal social systems, Trends in Ecology and Evolution 28 (2013) 541–551.

[4] J. J. St Clair, Z. T. Burns, E. M. Bettaney, M. B. Morrissey, B. Otis, T. B. Ryder, R. C. Fleischer, R. James, C. Rutz, Experimental resource pulses influence social-network dynamics and the potential for information flow in tool-using crows, Nature Communications 6 (2015) 1–8.

[5] A. Ilany, E. Akçay, Social inheritance can explain the structure of animal social networks, Nature Communications 7 (2016).

[6] R. K. Hamede, J. Bashford, H. McCallum, M. Jones, Contact networks in a wild Tasmanian devil (Sarcophilus harrisii) population: Using social network analysis to reveal seasonal variability in social behaviour and its implications for transmission of devil facial tumour disease, Ecology Letters 12 (2009) 1147–1157.

[7] J. A. Galbraith, M. C. Stanley, D. N. Jones, J. R. Beggs, Experimental feeding regime influences urban bird disease dynamics, Journal of Avian Biology 48 (2017) 700–713.

[8] K. E. Jones, N. G. Patel, M. A. Levy, A. Storeygard, D. Balk, J. L. Gittleman, P. Daszak, Global trends in emerging infectious diseases, Nature 451 (2008) 990–993.

[9] S. Davis, B. Abbasi, S. Shah, S. Telfer, M. Begon, Spatial analyses of wildlife contact networks, Journal of the Royal Society Interface 12 (2015).

[10] W. Ji, P. C. White, M. N. Clout, Contact rates between possums revealed by proximity data loggers, Journal of Applied Ecology 42 (2005) 595–604.

[11] J. A. Drewe, N. Weber, S. P. Carter, S. Bearhop, X. A. Harrison, S. R. Dall, R. A. McDonald, R. J. Delahay, Performance of proximity loggers in recording intra- and inter-species interactions: A laboratory and field-based validation study, PLoS ONE 7 (2012).

[12] E. Sheehy, C. Sutherland, C. O’Reilly, X. Lambin, The enemy of my enemy is my friend: Native pine marten recovery reverses the decline of the red squirrel by suppressing grey squirrel populations, Proceedings of the Royal Society B: Biological Sciences 285 (2018).

[13] D. P. Croft, S. K. Darden, T. W. Wey, Current directions in animal social networks, Current Opinion in Behavioral Sciences 12 (2016) 52–58.

[14] M. J. Greenlees, G. P. Brown, J. K. Webb, B. L. Phillips, R. Shine, Do invasive cane toads (Chaunus marinus) compete with Australian frogs (Cyclorana australis)?, Austral Ecology 32 (2007) 900–907.

[15] H. Leirs, Population ecology of Mastomys natalensis (Smith, 1834). Implications for rodent control in Africa, Ph.D. thesis, University of Antwerp, 1994.

[16] J. J. Reynolds, B. T. Hirsch, S. D. Gehrt, M. E. Craft, Raccoon contact networks predict seasonal susceptibility to rabies outbreaks and limitations of vaccination, Journal of Animal Ecology 84 (2015) 1720–1731.

[17] F. Maroto-Molina, J. Navarro-García, K. Príncipe-Aguirre, I. Gómez-Maqueda, J. E. Guerrero-Ginel, A. Garrido-Varo, D. C. Pérez-Marín, A low-cost IOT-based system to monitor the location of a whole herd, Sensors (Switzerland) 19 (2019).

[18] S. P. Ripperger, S. Stockmaier, G. G. Carter, Tracking sickness effects on social encounters via continuous proximity sensing in wild vampire bats, Behavioral Ecology 31 (2020) 1296–1302.

[19] M. Bohm, K. L. Palphramand, G. Newton-Cross, M. R. Hutchings, P. C. White, Dynamic interactions among badgers: implications for sociality and disease transmission, Journal of Animal Ecology (2008) 281–291.

[20] M. E. Craft, E. Volz, C. Packer, L. A. Meyers, Disease transmission in territorial populations: The small-world network of Serengeti lions, Journal of the Royal Society Interface 8 (2011) 776–786.

[21] R. Kays, M. C. Crofoot, W. Jetz, M. Wikelski, Terrestrial animal tracking as an eye on life and planet, Science 348 (2015) aaa2478.

[22] R. Walrath, T. R. Van Deelen, K. C. Vercauteren, Efficacy of proximity loggers for detection of contacts between maternal pairs of white-tailed deer, Wildlife Society Bulletin 35 (2011) 452–460.

[23] I. I. Levin, D. M. Zonana, J. M. Burt, R. J. Safran, Performance of encounternet tags: Field tests of miniaturized proximity loggers for use on small birds, PLoS ONE 10 (2015) 1–18.

[24] S. Ripperger, D. Josic, M. Hierold, A. Koelpin, R. Weigel, M. Hartmann, R. Page, F. Mayer, Automated proximity sensing in small vertebrates: Design of miniaturized sensor nodes and first field tests in bats, Ecology and Evolution 6 (2016) 2179–2189.

[25] B. Cassens, S. Ripperger, M. Hierold, F. Mayer, R. Kapitza, Automated Encounter Detection for Animal-Borne Sensor Nodes, EWSN ’17 Proceedings of the 2017 International Conference on Embedded Wireless Systems and Networks (2017) 20–22.

[26] N. Duda, T. Nowak, M. Hartmann, M. Schadhauser, B. Cassens, P. Wägemann, M. Nabeel, S. Ripperger, S. Herbst, K. Meyer-Wegener, E. Mayer, F. Dressler, W. Schröder-Preikschat, R. Kapitza, J. Robert, J. Thielecke, R. Weigel, A. Kölpin, Bats: Adaptive ultra low power sensor network for animal tracking, Sensors (Switzerland) 18 (2018).

[27] F. Dressler, M. Mutschlechner, M. Nabeel, J. Blobel, Ultra Low-Power Sensor Networks for Next Generation Wildlife Monitoring, 2019 11th International Conference on Communication Systems and Networks, COMSNETS 2019 2061 (2019) 44–48.

[28] E. D. Ayele, N. Meratnia, P. J. Havinga, Towards a new opportunistic iot network architecture for wildlife monitoring system, 2018 9th IFIP International Conference on New Technologies, Mobility and Security, NTMS 2018 - Proceedings 2018-Janua (2018) 1–5.

[29] M. Aernouts, R. Berkvens, K. Van Vlaenderen, M. Weyn, Sigfox and LoRaWAN datasets for fingerprint localization in large urban and rural areas, Data 3 (2018) 1–15.

[30] F. Lemic, V. Handziski, M. Aernouts, T. Janssen, R. Berkvens, A. Wolisz, J. Famaey, Regression-Based Estimation of Individual Errors in Fingerprinting Localization, IEEE Access 7 (2019) 33652–33664.

[31] M. Roeleke, T. Blohm, S. Kramer-Schadt, Y. Yovel, C. C. Voigt, Habitat use of bats in relation to wind turbines revealed by GPS tracking, Scientific Reports 6 (2016) 1–9.

[32] R. Berkvens, I. H. Olivares, S. Mercelis, L. Kirkpatrick, M. Weyn, Contact detection for social networking of small animals, Lecture Notes on Data Engineering and Communications Technologies 24 (2019) 405–414.

[33] S. P. Ripperger, G. G. Carter, R. A. Page Supervision, N. Duda, A. Koelpin, R. Weigel, M. Hartmann, T. Nowak, J. Thielecke, M. Schadhauser, J. Robert, S. Herbst, K. Meyer-Wegener, P. Wägemann, W. S. Preikschat, B. Cassens, R. Kapitza, F. Dressler, F. Mayer, Thinking small: Next-generation sensor networks close the size gap in vertebrate biologging, PLoS Biology 18 (2020) 1–25.

[34] Bluetooth Smart or Version 4.0+ of the Bluetooth specification, https://www.bluetooth.com/, 2010. Rev. 4.0.

[35] U. M. Qureshi, F. K. Shaikh, Z. Aziz, Z. Shah, A. Sheikh, E. Felemban, S. Qaisar, RF Path and Absorption Loss Estimation for Underwater Wireless Sensor Networks in Different Water Environments, Sensors 16 (2016).

[36] BGM111 Blue Gecko Bluetooth Module Data Sheet, Silicon Labs, 2018. Rev. 1.4.

[37] BGM121/BGM123 Blue Gecko Bluetooth SiP Module Data Sheet, Silicon Labs, 2018. Rev. 1.3.

[38] C. N. Ltd., Tag-connect,llc, http://www.tag-connect.com/, 2018.

[39] N. Semiconductors, nrf52 development kit product brief, http://infocenter.nordicsemi.com/pdf/nRF52_DK_PB_v2.0.pdf, 2018. Rev. 2.0.

[40] M. Ghamari, E. Villeneuve, C. Soltanpur, J. Khangosstar, B. Janko, R. S. Sherratt, W. Harwin, Detailed Examination of a Packet Collision Model for Bluetooth Low Energy Advertising Mode, IEEE Access 6 (2018) 46066–46073.

[41] B. Borremans, N. K. Hughes, J. Reijniers, V. Sluydts, A. A. Katakweba, L. S. Mulungu, C. A. Sabuni, R. H. Makundi, H. Leirs, Happily together forever: Temporal variation in spatial patterns and complete lack of territoriality in a promiscuous rodent, Population Ecology 56 (2014) 109–118.

[42] C. Rutz, M. B. Morrissey, Z. T. Burns, J. Burt, B. Otis, J. J. St Clair, R. James, Calibrating animal-borne proximity loggers, Methods in Ecology and Evolution 6 (2015) 656–667.

[43] R. James, D. P. Croft, J. Krause, Potential banana skins in animal social network analysis, Behavioral Ecology and Sociobiology 63 (2009) 989–997.

[44] C. Rutz, Z. T. Burns, R. James, S. M. Ismar, J. Burt, B. Otis, J. Bowen, J. J. St Clair, Automated mapping of social networks in wild birds, Current Biology 22 (2012) R669–R671.

[45] M. K. Marsh, S. R. McLeod, M. R. Hutchings, P. C. White, Use of proximity loggers and network analysis to quantify social interactions in free-ranging wild rabbit populations, Wildlife Research 38 (2011) 1–12.

[46] C. C. Wilmers, B. Nickel, C. M. Bryce, J. A. Smith, R. E. Wheat, V. Yovovich, M. Hebblewhite, The golden age of bio-logging: How animal-borne sensors are advancing the frontiers of ecology, Ecology 96 (2015) 1741–1753.

